# In vivo Ontogeny of human forebrain neural progenitor cell grafts in adult rats: an immunohistological study

**DOI:** 10.1101/2023.05.15.539796

**Authors:** Chunhua Liu, Aiping Lin, Qi Xing, Di Zhang, Wei Meng, Rui Wu, Heng Shi, Wenhao Huang, Xiaofen Huang, Dajiang Qin, Xiaoyun Wang, Xiaofen Zhong, Yiping Guo

**Affiliations:** Guangdong Provincial Key Laboratory of Stem Cell and Regenerative Medicine, CAS Key Laboratory of Regenerative Biology, Guangzhou Institutes of Biomedicine and Health, Chinese Academy of Sciences, Guangzhou, 510530, Guangdong Province, China; The Fifth Affiliated Hospital of Guangzhou Medical University, Guangzhou, 510700, China; Biotherapy Centre, The Third Affiliated Hospital, Sun Yat-sen University, Guangzhou, 510630, China; Guangdong Work Injury Rehabilitation Center, Guangzhou, 510440, China; Department of Rehabilitation, The Third Affiliated Hospital, Sun Yat-sen University, Guangzhou, 510630, China; Center for Cell Lineage and Development, Guangzhou Institutes of Biomedicine and Health, Chinese Academy of Sciences, Guangzhou, 510530, China; Department of Neuroscience, City University of Hong Kong, Kowloon Tong, Kowloon, Hong Kong SAR; Guangzhou Regenerative Medicine and Health GuangDong Laboratory (GRMH-GDL), Guangzhou, 510005, China

**Keywords:** Neural progenitor cell, Cell fate tracing, Transplantation, Migration, Ontogeny

## Abstract

A thorough understanding of the cell behaviors of the human neural grafts is fundamental to exploit them to achieve cell therapy for recovering brain functions. Here by using immunohistological staining, we trace the cell fate of the intrastriatal human neural progenitor cell (NPC) grafts up to 9 months in adult rats, with multiple examining time points to provide a unified working time frame for future transplantation study. Lots of Nestin+/Sox2+ human cells continuously migrate along the white matter tracts into distal brain parenchyma even long time after transplantation, providing a potential for curing diffuse brain damage. Further analysis reveals a significant heterogeneity of the long-term sustained neural stem cells (NSC)/NPCs that progressing throughout different stages, mimicking the neural development of human forebrain. More importantly, the initial GFAP expression in human grafts marks the NSC progression instead of terminal astrocyte differentiation. The distally migrating human cells continuously show the capability to produce new neurons, albeit at a low efficiency in the intact brain. Further investigations in neural disease models are needed. Such study would benefit neural cell therapy with regarding to the optimization of the transplantation strategy and choosing of acting mode by neural grafts (e.g. via cell replacement or ***ex vivo*** gene therapy).

## Introduction

Human neural stem cells hold the promise for neurological disorders and irreversible brain injuries. Thus a thorough understanding of the cell behaviors of the human neural grafts is important, especially their cell fates in the adult brain, including their survival, proliferation, migration, differentiation, maturation and integration to the host neural circuit, as well as the pruning of redundant neural progenitor cells (NPCs) or their derivatives. These knowledges would facilitate the use of the human neural stem cells (NSCs)/NPCs to achieve cell therapy. It can be realized via cell replacement or targeted delivery of neuroprotective and regenerative protein(s) for ***ex vivo*** gene therapy, after optimization of the transplantation strategy (e.g. the transplantation site, timing, proper NPC passage selected or what gene modification used) to maximize benefits and minimize risks. To fulfill this perspective, extensive studies have been carried out by transplanting human NPCs in intact [1-4] or damaged brain [5-10], in the adult, neonatal [11] or fetal [12] rodents, as well as in the non-human primate disease models [13, 14]. However, most of the studies only examined one or 2-3 time point(s) after transplantation, thus failed to provide a unified time course of the cell fates of the human neural grafts, leading to contradictionary conclusions. More importantly, the expression of cell markers is a dynamic process, with a restricted spatiotemporal control. For example, ***in vitro*** neural induction and differentiation system has used GFAP as a reliable astrocyte marker, so do many ***in vivo*** studies [15-17]. However, a totally different scenario shows up when examining the GFAP expression during development. GFAP is expressed in the radial glial cells (RGC) and immature astrocytes, and retains in the SVZ radial glia-like cells of adults, but is repressed in mature astrocytes [18] (for review, see [19, 20]). The geographic expression of GFAP is another important factor to define the cell fate of the GFAP+ cells. The initial expression of GFAP in neurogenic niche (e.g. SVZ) and in the non-neurogenic regions (e.g. the immediate zone and the cortical plate) occurs almost at the same time. It marks the late RGC in SVZ while labels the earliest astrocytes or the astrocyte precursors in the cortical plate during development.

In the present study, we traced the cell fate of the human ESC-derived forebrain committed NPC grafts up to 9-month long after transplantation into the intact brain of rats, with multiple examining time points including 4 days post-transplantation (dpt), 9/14 dpt, 3/4 weeks post-transplantation (wpt), 6 wpt, 8 wpt, 12/13 wpt, 16/17 wpt, 24/26 wpt and 38/41 wpt. To avoid the lateral ventricles and SVZ where adult NSCs resident, we transplanted human NPCs into the dorsolateral striatum. After a short adaptive phase following transplantation, human cells resumed to proliferate and differentiate, building up a human graft core locally within 3 weeks [4]. Then, human cells gradually spread away into the host territory in a radial diffusion-like way, forming a graft colony, where the human cells and the local host cells showed a salt-and-pepper distribution. As previously reported [4, 21], these human cells showed intermingled expressions of Nestin, DCX and TUJ1 soon after transplantation. Human cells also migrated widely along the white matter tracts into areas far beyond the transplantation site as others documented [1, 2]. Unexpectedly, at 41 wpt, we still can see lots of Nestin+ human cells scattered in the host brain parenchyma without rosette structures or a specialized stem cell “niche” to support them, which might promise a long-term benefit for regeneration or delivery of targeted medicine, but also relate to the overgrowth risk and time efficiency of neural stem cell therapy, which further warranting the present study.

## Results

### Long-term sustaining of human NPCs after transplantation

Nestin and Sox2 have been widely used as markers of neural stem/ progenitor cells that progress through different stages and vanish upon differentiation. By taking advantages of the antibody against human specific Nestin (hNestin), we first examined Nestin expression in human cells at different assessment time points after transplantation to evaluate the in vivo maintenance of human grafts. Typical neural rosette structures consisting of slowly proliferating neural stem cells manifested within 2-3 wpt when forebrain NPCs transplanted [4]. As expected, hNestin was widely expressed in the neural rosette, as well as in quite a few non-rosette cells at 2-8wpt, whereby the Nestin+ ribbons spread throughout the whole graft colony (**Figure 1A**). The hNestin+ cells were also among the radially migrating population, where their morphology became distinguishable. The majority of hNestin+ cells were bipolar within 12 wpt, which gradually protruded more processes and became star-shaped (**Figure 1A**). In some more distal areas, only numerous hNestin+ ribbons without visible soma were seen, indicative of wide diffusion or migration of the long-term sustained human NPCs. The hNestin+ fibers intermingled to each other, making it difficult to recognize individual cell bodies and thus difficult for quantification.

**Figure 1.**
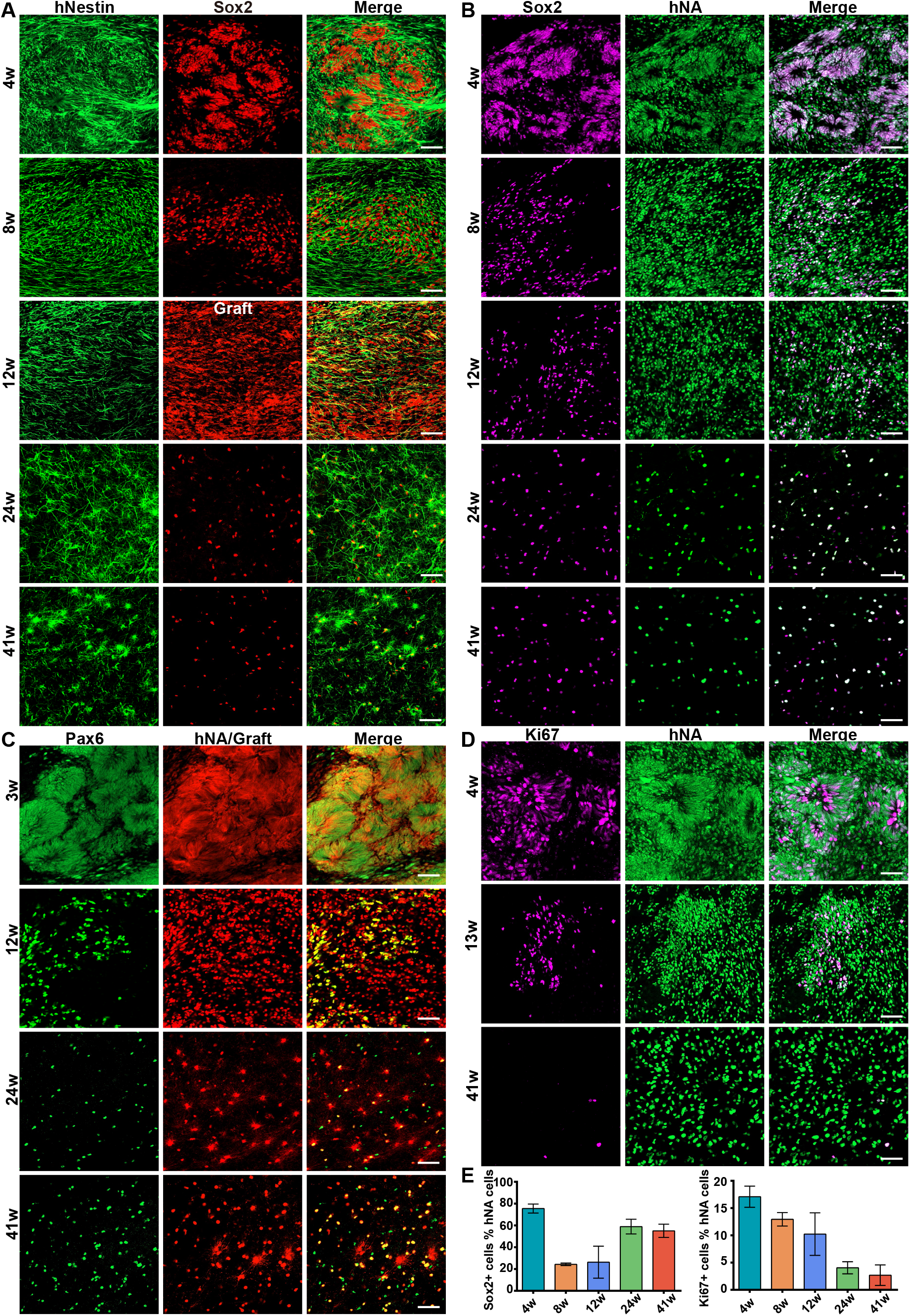
Expressions of neural stem cell markers (Nestin, Sox2 and Pax6) and proliferation marker Ki67 by human neural grafts. **A)** human specific Nestin and Sox2, *Graft* means human graft fluorescent reporter in all figures; **B)** Sox2 and human nucleus-specific antibody hNA; **C)** Pax6 and hNA alone or hNA combined with graft reporter; **D)** Ki67 and hNA; **E)** (left) quantification of Sox2+ cells among human cells; (right) quantification of Ki67+ cells among human cells. Scale bar: 50μm, unless otherwise stated.

We then used the nucleus-located Sox2 to further quantify the human NPCs. Double staining against hNestin and Sox2 showed that the majority of hNestin+ cells clearly expressed Sox2, and nearly all Sox2+ expressed hNestin (**Figure1A**). At 4 wpt, similar to Nestin, most of human cells (75.5±4.1%) showed Sox2 expression. Concomitant with the disappearance of rosettes at 8 wpt, Sox2 expression decreased quickly in scattered cells upon their proceedings to neuronal differentiation, while retained a high level in the cell clusters (**Figure 1B**). Surprisingly, at late time points, the Sox2+ human cells took a relatively higher proportion (58.9±6.8 % at 24wpt; 55.0±6.1 % at 38/41 wpt) among the scattered cells when the early cell clusters had largely dispersed (**Figure 1B**). Pax6 is an important marker of human neuroepithelial cells as well as RGCs. At 4wpt, Pax6 was expressed in the rosette neural stem cells (R-NSC), while many peripheral non-rosette cells decreased its expression. Similar to Sox2, Pax6+ human cells again accounted for a higher proportion among the scattered cells at later time points (**Figure 1C**).

To measure the proliferation capability of the Nestin+/Sox2+ human cells, we did Ki67 staining (**Figure 1D**). Ki67+ human cells were detected mainly within the graft colony, which decreased sharply over time from 17.1±1.9% at 4 wpt to a very low level of 4.0±1.1% at 24 wpt and 2.7±1.9% at 38/41 wpt, similar to that of the adult NSCs in the neurogenic niche [22]. Ki67 was also expressed in distally migrating human cells at all time points, but in an even lower ratio. These results greatly alleviated the concerns about the overgrowth risk and tumorigenicity of the human NPC grafts.

### Time-course protein expression profile of RGC markers

Previous studies have revealed a highly heterogeneity of neural stem cells and neural precursor cells during human embryonic development [23, 24]. To further figure out whether heterogeneity is also gradually acquired by the long-term sustained human NPC grafts and to trace their ontogeny, we performed detailed immunostainings against a series of stage-specific or stage-preferred markers, including Pax6/Sox2/Nestin for Pan-NSCs (see above), BLBP/Vimentin/Hes5/GFAP for RGCs, TBR2 for immediate progenitor cells (IPCs), GFAP/AQP4/ glutamine synthetase (GS)/ S100β for terminally differentiated astrocytes, DCX/ASCL1 for early neuronal progenitor cells, HuD/NeuN for neurons, NG2/MBP for oligodendrocyte lineage.

BLBP is among the earliest RGC markers beside RC1/RC2 antigens encoded by Nestin. It is transiently expressed in nearly all early ventricular RGCs during corticogenesis and required for the establishment of the radial glial fiber system [25]. Our data, however, showed that BLBP was detected sparkling only in a subgroup of human cells early at 9 dpt and then increased gradually to a level of ∼30-40% at 24-41 wpt. Surprisingly, It was virtually absent from the R-NSCs (**Figure 2A**). Since BLBP has been suggested to be induced in RGCs by neurons attaching to them [25], the sparse BLBP expression might imply that most of our human RGCs should not have neuroblasts crawling along them and thus did not play a major role in providing such a scaffold for radial migration. Echoing with this, BLBP stained the nucleus as well as the cytoplasm, lighting-up their cell soma clearly without radial BLBP+/RC2+ processes (**Figure 2A**), which, however, typically seen during corticogenesis.

**Figure 2.**
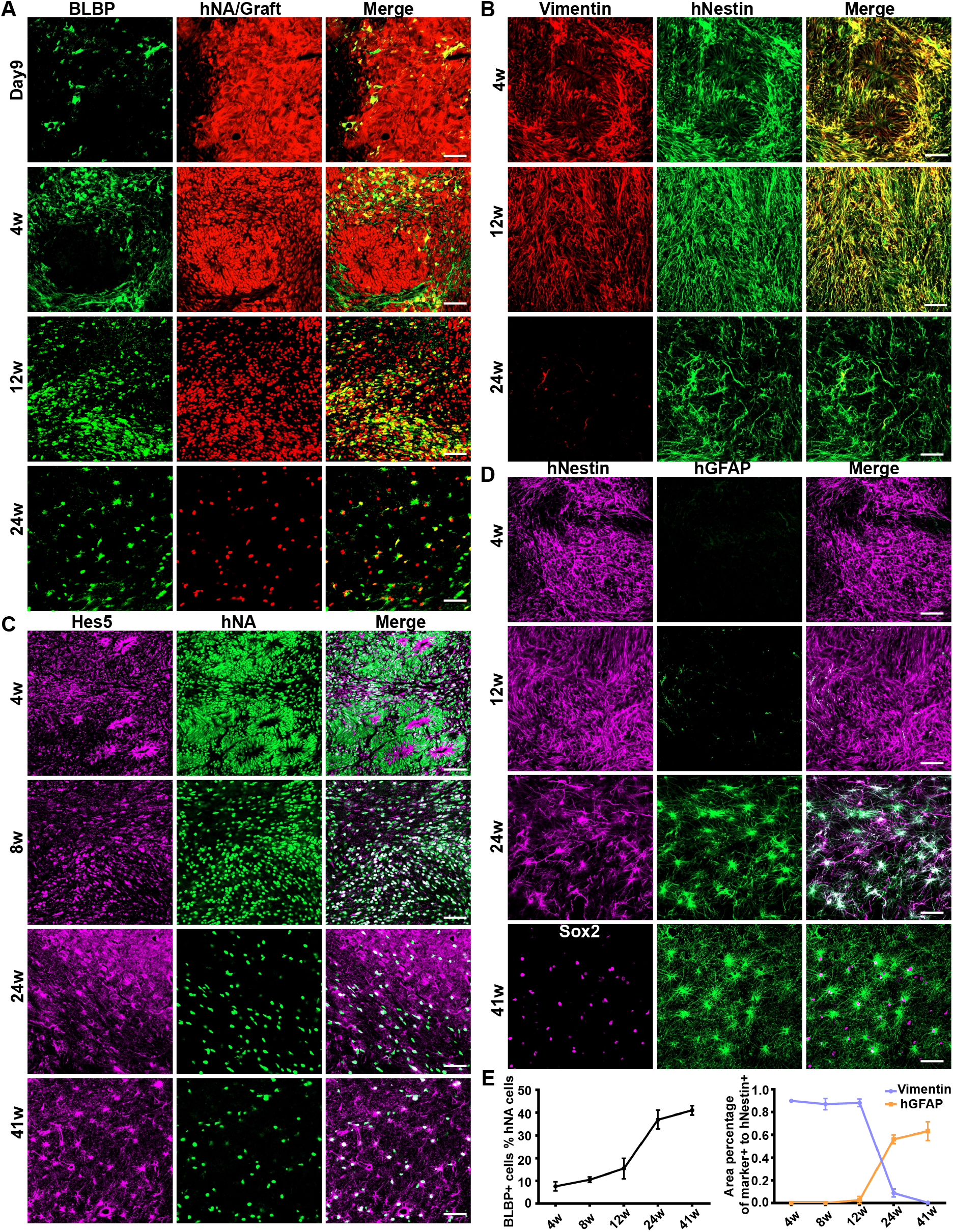
Expressions of RGC markers (BLBP, Vimentin, Hes5 and GFAP) by human neural grafts. **A)** BLBP and hNA or graft reporter; **B)** Hes5 and hNA; **C)** Vimentin and human Nestin; **D)** human specific GFAP and human Nestin or Sox2; **E)** (left) quantification of BLBP+ cells among human cells; (right) quantification of Vimentin+ and GFAP+ cells among human Nestin+ cells. Scale bar: 50μm.

Vimentin is an intermediate filament protein first expressed in early RGCs, and serves as a core marker of RGCs, participating in several key cellular events like the epithelial-mesenchymal transition (EMT) and migration. Strong Vimentin expression was detected early at 3-4 wpt. Different from BLBP, Vimentin expression is virtually seen in all R-NSCs. It maintained a high level till 16wpt, and then followed by a gradual decrease to a quite low level at 41 wpt (**Figure 2B**), indicating a waning EMT and radial diffusion in the human graft colony at late stages.

Hes5 is a nuclear protein and has been suggested to play a key role in maintaining of neural stem cells throughout the development [26, 27]. At neural rosette stage, Hes5 was moderately expressed in all NSCs, surprisingly with a relative high intensity in the apical cytosol, demarcating the central lumen (**Figure 2C**). After disappearance of rosettes, many human cells still showed a high Hes5 expression level, with a non-polarized cytosol staining. Nevertheless, we also saw nucleus-restricted localization of Hes5 in some samples (one example at 8 wpt shown in **Figure 2C**). The reason for the different subcellular distribution pattern of Hes5 is yet unknown.

GFAP is another intermediate filament protein which has been shown to replace the Vimentin in the late RGCs in rodents at the end of gestation that would give rise to astrocytes perinatally [19]. In the developing human and non-human primate brain, however, GFAP is expressed by RGCs early at the first trimester of gestation, thus also suggested as a marker of embryonic RGCs in human [28]. By using human specific GFAP (hGFAP) antibody, we found that hGFAP did not manifest until 12 wpt, and then increased rapidly, coincidentally with the decrease of Vimentin (**Figure 2D**). Double staining showed that nearly all hGFAP+ cells at early stage co-expressed Nestin or Sox2, suggestive of their NSC identity. GFAP+ cells among the hNestin+ cells quickly increased from 2.7±3.2% at 12 wpt to 56.2±3.9% at 24 wpt and 63.3±8.2% at 38/41 wpt, indicating that GFAP+/Nestin+ human cells accounted for a large proportion among the NSC-like cells at late stage.

### Initial GFAP expression in human transplants marking the NSC progression instead of terminal astrocyte differentiation

Given that GFAP is also a marker of immature post-mitotic astrocytes during development, by using several previously well-established astrocyte markers, S100β, GS and AQP4, we further examined whether some of hGFAP+ cells might have differentiated into the terminal astrocytes other than the late RGCs.

As S100β is not expressed by RGCs, NPCs and astrocyte progenitors, the co-expression of GFAP and S100β has been used as the “golden standard” of terminally differentiated astrocyte ***in vitro*** and ***in vivo*** [15-17]. No or very few human cells were positive in S100β staining before 19 wpt (including 19 wpt, **Figure 3A**). It was until 24 wpt that we could detect a few clear S100β+ human cells, most of which only showed weak S100β expression that confined in the perinuclear area, suggestive of the initial differentiation of human astrocytes (**Figure 3A**). Also from this time point and thereafter, very few hGFAP+/S100β+ cells were unveiled (**Figure 3B**). The ratio of the S100β+ cells among hGFAP+ human cells increased at 38/41 wpt, and some of them already showed a high S100β expression level (**Figure 3B-C**). We failed to find S100β+/hNestin+ double positive human cells at all time points, again implying that the onset of S100β expression marked the initial differentiation of human astrocytes from our NPC grafts.

**Figure 3.**
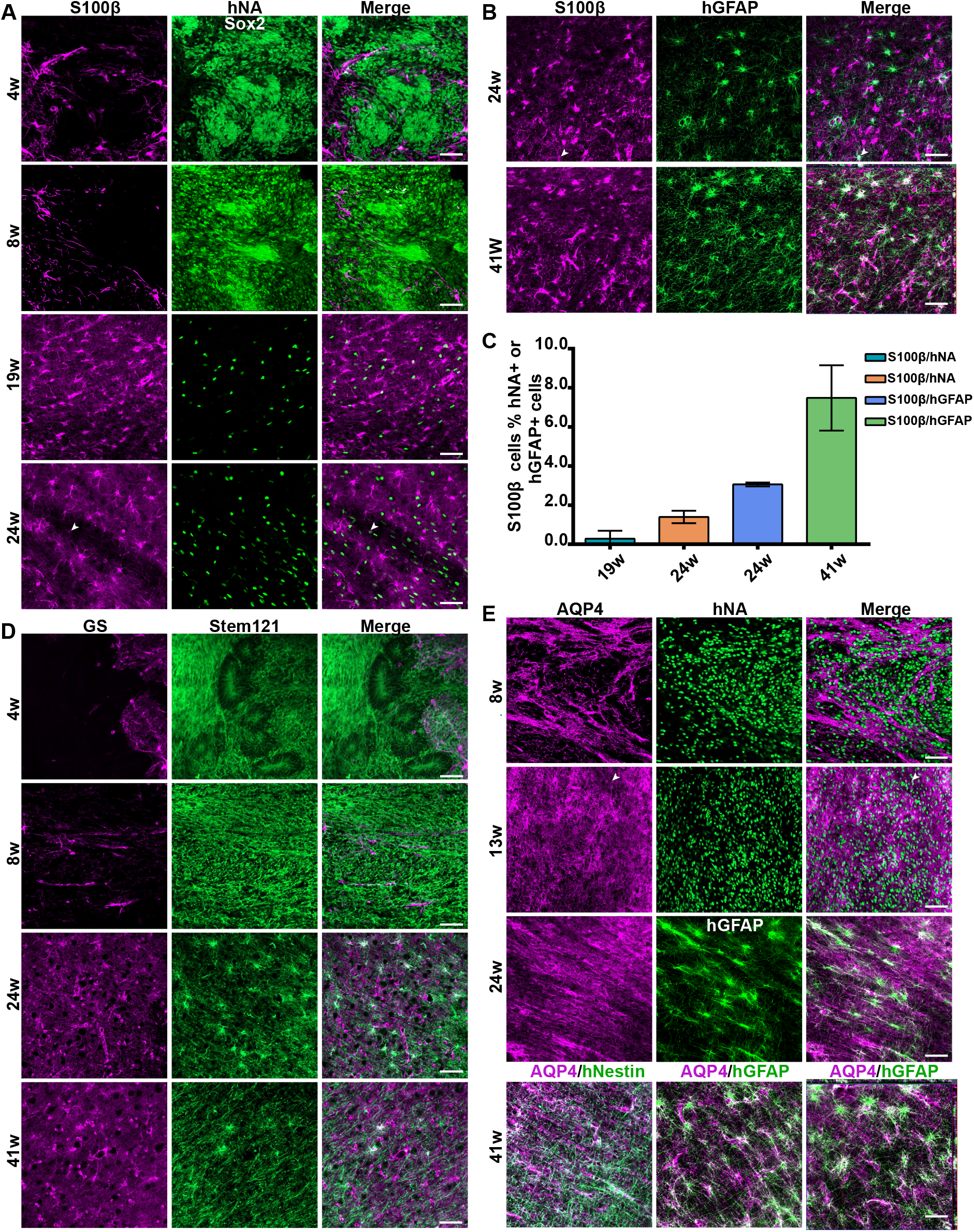
Expressions of astrocyte markers (S100β, GS and AQP4) by human neural grafts. **A)** S100β and hNA, *arrow heads* indicating the S100β+/hNA+ double positive cells with weak S100β expression; **B)** S100β and hGFAP+, *arrow heads* indicating the S100β+/hGFAP+ double positive cells; C) quantification of **S100β**+ cells among human cells and human GFAP+ cells; D) GS and human specific cytoplasmic antibody Stem121; **E)** AQP4 and hNA, hNestin or hGFAP. Scale bar: 50μm.

GS has also been reliably used as an astrocyte marker in many studies [29]. Immunostaining against GS showed similar results to S100β staining that human cells rarely expressed GS at any time point examined, although a couple of GS+ cells were observed at 24 wpt and onward (**Figure 3D**). Interestingly, GS+ immunoreactivity from the host cells invaded the graft core much earlier than the host GFAP, possibly from endothelial cells, consistent with a reported role of GS in angiogenesis [30]. Similar phenomena were also observed in S100β and AQP4 staining, both of which are also expressed in endothelial cells or perivascular membranes of the brain, particularly in response to an intervening event like stress [31, 32].

AQP4 is the marker of astrocytic end-feet in adults, where it serves as influx routes for water. Very few AQP4 positive-like human cells could be seen at the border of the grafts at 13 wpt and 16 wpt, but they never showed an end-feet structure (**Figure 3E)**. At 24 wpt and onward, some radial hGFAP+ human cells had a weak AQP4 expression, but still without end-feet structures, suggesting a developmental transition from a neural stem/progenitor cell identity to immature astrocytes. Although more hGFAP and AQP4 immunoreactivities were found to be co-localized at 38/41 wpt, most of AQP4 signals were actually derived from the host vascular structures, suggestive of glial-vascular units being assembled between donor and host upon early human astrocyte differentiation [33]. The hGFAP+/AQP4+ double positive cells only accounted for a small sub-population of hGFAP+ human cells. Similar to S100β, no or very few AQP4+/hNestin+ double positive human cells were found at all time points. While many hGFAP+ processes ran along the host AQP4+ ribbons, the hNestin+ fibers preferred to extend in different directions (**Figure 3E, Bottom**).

We further stained NG2/MBP for oligodendrocyte lineage differentiation, to provide more evidences to define the progression stage where these long-term sustained GFAP+ human cells situate, and found no or very few NG2+ and MBP+ human cells (0-1 per slice).

Collectively, our data demonstrated that the initial GFAP expression in human NPC graft marked the NSC progression instead of terminally astrocyte differentiation, and most of GFAP+ cells stayed in a “stem” or “transitional” state.

### Migration of Nestin+ and DCX+ human cells

DCX+ and Nestin+ human cells can be found immediately following transplantation. DCX is a migrating neuroblast marker. DCX+ cells have a typical leading protrusion or fiber, which is important to initiate the migration. Besides DCX+ cells, we observed that Nestin+ cells also protruded leading fibers. Both of them migrated into the host territory in a radial diffusion-like way and resided locally to form a graft colony. Moreover, their leading fibers could also reach out the white matters, e.g. subcortical corpus callosum and external capsule, and guided the DCX+ and Nestin+ human cells to enter the host white matter tracts. We further confirmed these migrating human Nestin+ cells are neural stem cells by co-staining with Sox2 and Pax6 (**Figure 4A**).

**Figure 4.**
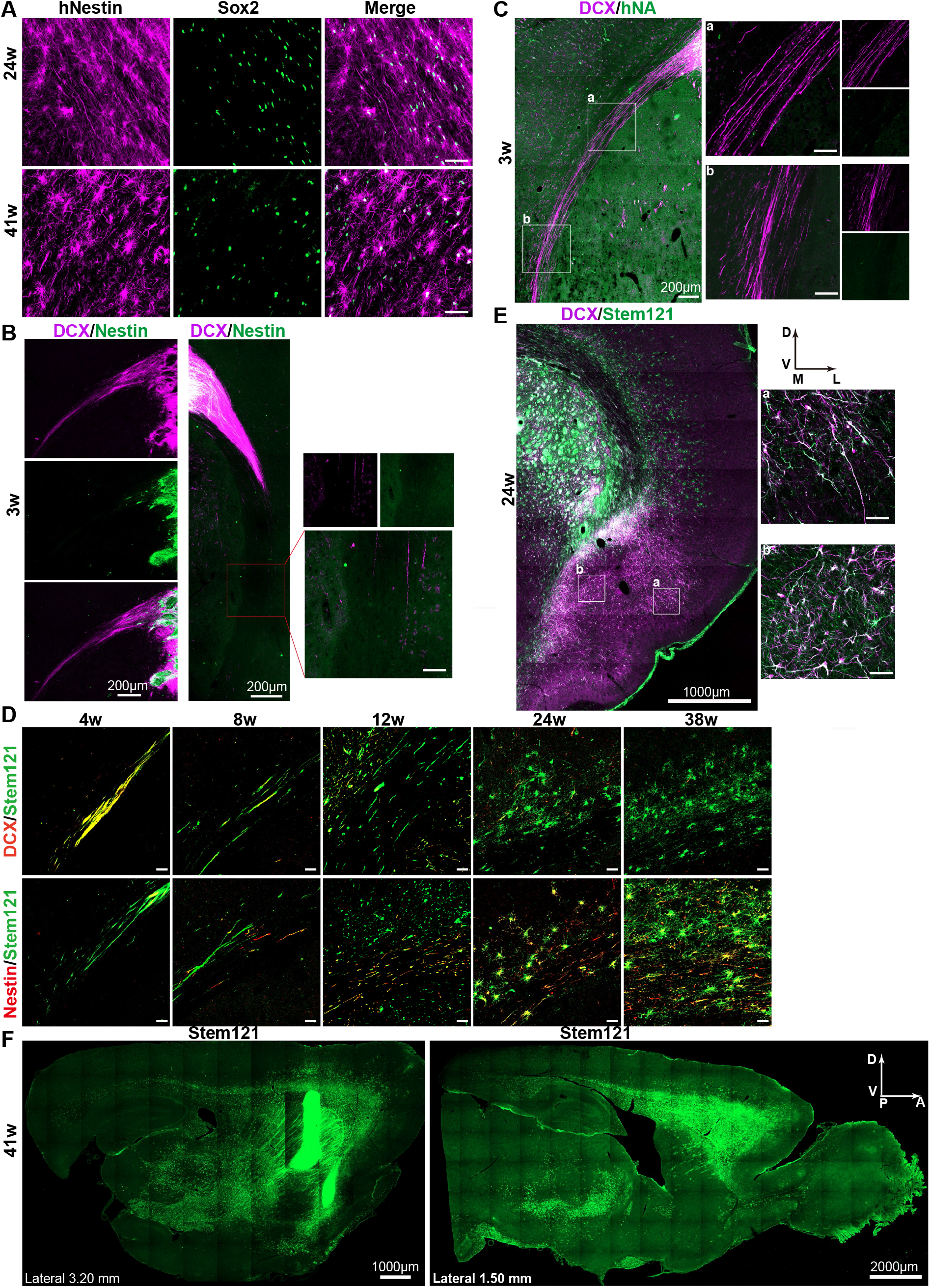
Expressions of Nestin and DCX by migratory human neural cells. **A)** human Nestin and Sox2; **B)** DCX and human Nestin at early stage, showing that DCX leading fibers extend far beyond the Nestin fibers; **C)** DCX and hNA expression in the migration pathway, showing that the human nuclei lagged far behind the DCX fibers; **D)** human specific cytoplasmic antibody Stem121 and DCX or human Nestin: long DCX fibers were only seen at early stages, bipolar DCX+ cells populated at 8 wpt and disappeared at 12 wpt from the migration pathway, while bipolar Nestin+ cells appeared at 8 wpt and populated at 12 wpt, then stellate Nestin+ cells gradually accounted for the majority. **E)** DCX and Stem121 at the distal temporal lobe at late stages; **F)** wide migration of human cells into thalamus, frontal lobe, olfactory bulb and posterior regions at sagittal plates. Scale bar: 50μm.

DCX+ fibers extended much faster than the Nestin+ fibers in both radial and tangential pathway (**Figure 4B**). Long DCX fibers were often seen to protrude and extend bidirectionally along the subcortical white matter tracts within a couple of weeks in coronal plane. By co-staining with hNA, we found the cell body of these fibers greatly lagged behind at 3 wpt, supporting the concept of leading fiber (**Figure 4C**). DCX+ cell bodies normally showed up in the white matters after that. DCX+ cells travelled possibly in a well-known manner that the leading fibers would retract from the lagged cell rear, pulling the cell body to move forward along the white matter fibers. DCX+ fibers reached out the ventrolateral end of external capsule in the ipsilateral hemisphere within 12 weeks (**Figure 4D, up**). Bipolar Nestin+ cells with well-recognized hNA+ nucleus and retracted leading fiber normally showed up in the corpus callosum at 8 wpt, populated in the whole path at 12 wpt, gradually protruded more processes and showed a star-shape at 16 wpt, with the same timeline of the radially diffused Nestin+ cells in the graft colony (**Figure 4D, down**). At 3-4 mpt, almost all long DCX+ fibers and DCX+ cells disappeared from the subcortical migration stream except a few remaining evidenced in the distal end (**Figure 4E)**. These data implied that Nestin+ cells could migrate independently on the DCX+ fibers, and the distal DCX+ cells detected at late time points (e.g. 24-41 wpt) might derive from the migrating Nestin+ cells.

With the marching of human cells along the white matter tracts, DCX+ cells were continuously migrating away into the overlying cortical layers in a radial diffusion manner from the very beginning, so were the Nestin+ cells. Interestingly, the claustrum and entorhinal cortex located at the distal end of the external capsule in the temporal lobe were among the most popular destinations (**Figure 4E)**, suggesting that the majority of migrating human cells in the subcortical white matter tracts would keep going until the very end of the migration path and then diffuse into the brain parenchyma there in intact brain. It is intriguing to know whether and to what extent they can be recruited by a midway or distal brain injury.

The human cells also migrated caudally and rostrally along the subcortical white matter tracts. They also ran into other white matter fibers sequentially, e.g. the internal capsule and anterior commissure, as well as corticostriatal projections, forming a continuum with the subcortical corpus callosum and external capsule, whereas very few of them would enter the SVZ-RMS-OB pathway. Human cells can go through these fibers to reach the thalamus, temporal lobe and olfactory bulb, as well as the ipsilateral sensorimotor cortex (that projecting to dorsolateral striatum), respectively (**Figure 4F**).

### Continuous differentiation of neuron-committed cells by the long-term sustained human NPCs

Great consensus has been reached that the grafted NPCs can produce neurons locally, just as we previously reported [4, 10, 21]. The question is whether the long-term sustained human NPCs were capable of generating neurons continuously. At all time points, we observed the intermingled distribution pattern of Nesin+ and DCX+ cells within the graft colony (**Figure 5A**). Notably, at 41 wpt, long after the disappearance of DCX+ cells from the migration stream, we could still see lots of DCX+ cells in the distal areas, suggesting that the migrating Nestin+ cells might give rise to these distal neuroblasts (**Figure 4E**). DCX+ neuroblasts would proceed to neurons, in which DCX was transiently retained.

**Figure 5.**
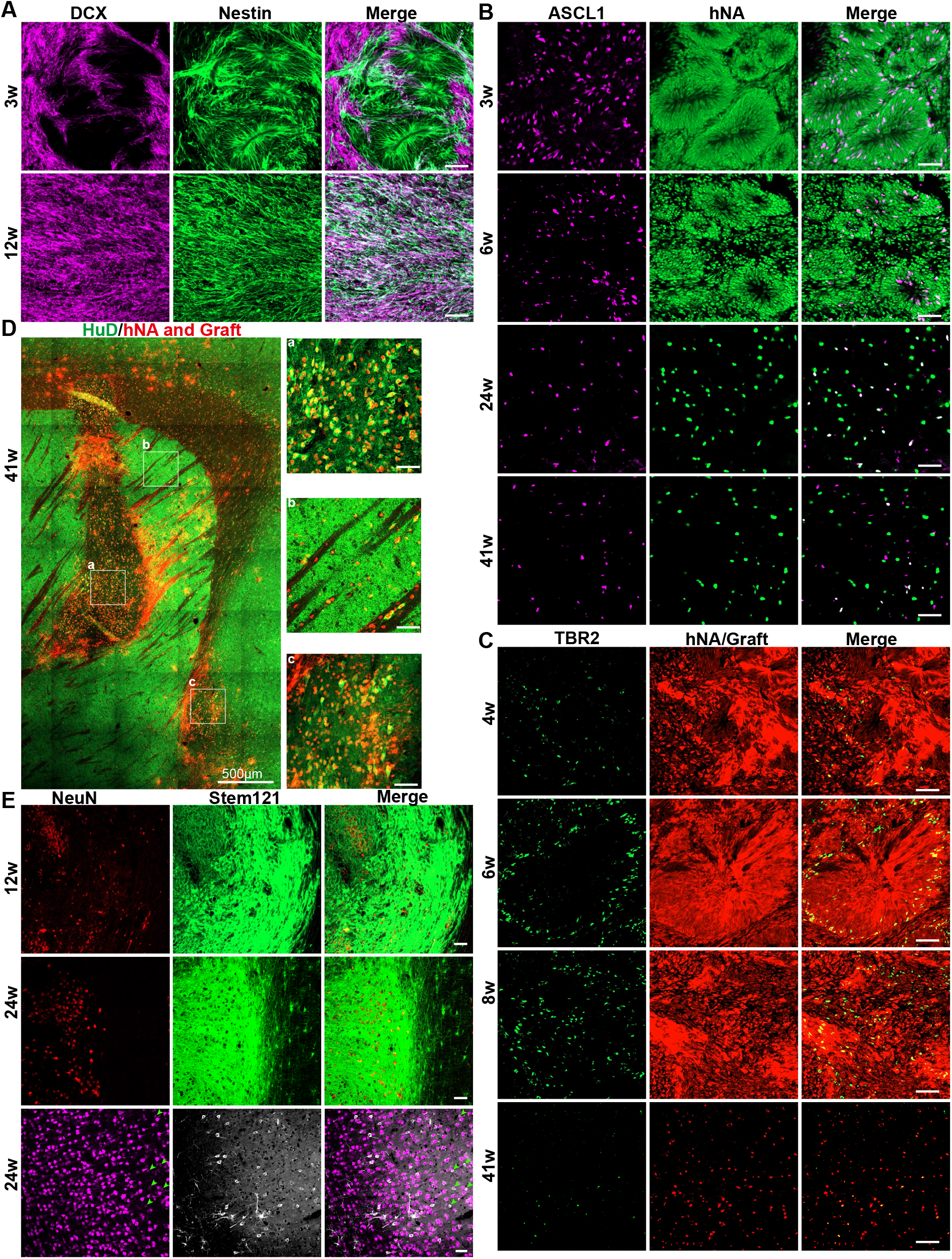
Expressions of pro-neuronal markers (DCX, ASCL1, TBR2, HuD and NeuN) by human neural grafts. **A)** the intermingled distribution of Nesin+ and DCX+ cells within the graft colony; **B)** ASCL1 and hNA; **C)** TBR2 and hNA or graft reporter; **D)** HuD and hNA combined with graft reporter: ***a,b,c*** respectively showing the graft colony, migration pathway and distal temporal area. **E)** NeuN and human Stem121: *TOP* and *Middle* panels showing the graft colony, *Bottom* showing the distal area, *arrowheads* indicating human cells of weak NeuN+ and Stem121+ immunoreactivities. Scale bar: 50μm.

ASCL1 is one of the earliest and driving markers for neuron-committed cell fate differentiation. ASCL1 was detected in both the rosettes and outside regions, with a comparable proportion (**Figure 5B**). After the disappearance of rosette structure, ASCL1 continued to express in a small subgroup of human cells (**Figure 5B**). However, we failed to detect double staining of ASCL1 and GFAP at any time point.

TBR2 is specifically expressed in the intermediate progenitor cells (IPCs) of the developing cerebral cortex, marking the manifestation of SVZ. TBR2 was seen in the peripheral region of the rosette, consistent with the ***in vitro*** data in other studies. After the disappearance of rosette structures, TBR2+ cells gradually decreased to a low but detectable level until 41 wpt (**Figure 5C**), implying that the long-term sustained Nestin+/GFAP+ NSCs could continuously give rise to neurons at least partially via the IPC indirect differentiation trajectory.

Collectively, all these data demonstrated that the long-term sustained human NPCs could continuously differentiate into neuron-committed cells. To further trace their cell fates, we performed immunostaining against more neuronal markers, i.e. HuD and NeuN. HuD is one of the earliest neuronal markers and expresses in both early and late neurons. HuD+ human cells were found mainly the graft colony, but also in the migration pathway and the overlying cerebral cortex, e.g. the distal claustrum and entorhinal cortex (**Figure 5D**). However, most of the distal HuD+ human cells failed to further mature into healthy NeuN+ neurons. NeuN+ human cells were collected in clusters at the graft colony, but only few of them were seen in the distal areas (**Figure 5E**). The distal NeuN+ human cells always had a weak NeuN immunoreactivity when compared with the host neurons and the human neurons at the colony as well. In the overlying cerebral cortex, human neurons that distal to migration pathway (indicated by arrowheads in **Figure 5E, Bottom/***right side*) showed a circular shape with fewer protrusions than the proximal ones (**Bottom***/Left side*), seemingly under degenerating. Given that there was no space for the distally-invaded and dispersed human neurons to fulfill their function in the intact cerebral cortex, they were supposedly removed via a clearing mechanism to maintain the homeostasis of the brain. However, we failed to detect significant TUNEL immunoreactivity, possibly due to the rapid clearance of cell debris, which has been estimated to occur within as little as 0.5 hours [34]. It is intriguing to know whether and to what extent the DCX+ human cells could become mature neurons and survive extensively when they were recruited by brain injury.

## Discussion

By using immunohistological stainings, the present study traced the cell fates and the migration behaviors of the intrastriatal human NPCs up to 9-month long in adult rats, with multiple examining time points, and provided a unified working time frame for future transplantation study and clinical application. The major finding incudes, 1) besides the locally settled down cells, human NPCs would enter into and migrated extensively along the white matter tracts into the distal brain parenchyma, in an identity of DCX+ transiently or Nestin+ cell persistently; 2) the long-term sustained human NSCs/NPCs were significantly heterogeneous that progressing throughout different stages, indicated by different combinations of expressed markers; 3) the initial GFAP expression in human grafts marks the NSC progression instead of terminal astrocyte differentiation, which mimicking the neural development, quite different from the ***in vitro*** study; 4) the long-term sustained human NPC grafts showed the capability to produce new neurons albeit at low efficiency more than half a year later, but their long-term survival and maturation in the intact brain remain doubt.

### The long-distance migration and maintenance of human NPC grafts holds the promise for curing diffuse brain damage

So far, for cell therapy, two strategies were used to select transplantation site. The first choice is to transplant the cells directly into the lesion sites [5, 8, 10], the neighboring areas [7] or in the denervated areas, e.g. the striatum as the transplantation site in Parkinson Disease model [35, 36]. These studies have showed that the locally injected NPCs can rebuild the damaged neural circuit and promote function recovery in rodents. Alternatively, cells were transplanted systemically/distally, e.g. intra-arterially, intravenously, intraventricularly, subarachnoidally or intranasally, due to the inaccessibility of the targeted region or widespread damage sites, and the small surgery preferred as well. For neuronal replacement therapy, the therapeutic efficacy of this strategy depends heavily on the capability of targeted migration of the transplanted cells. Endogenous adult NSCs from SVZ migrate along the SVZ-RMS-OB route in the form of neuroblasts. Although Nestin+ NSCs have been shown within the route, they are considered as the local residents, not derived from the SVZ [37]. The NSCs from SVZ and from the RMS have different differentiation potency: the former differentiates into new neuron in OB, while the latter generates significantly more oligodendrocytes [37]. Following brain injury, the SVZ neuroblasts can also migrate to the lesion site, especially for the striatum injury, where they can differentiate into neurons and to some extent, contribute to the function recovery in rodents. Faiz and colleagues, by using lineage tracing strategy with Nestin-Cre mouse lines, showed that SVZ multipotent NSCs leaved the SVZ niche, migrated to the site of injury following stroke, while retained their ability to form multipotent neural spheres ***in vitro***, and but eventually gave rise to reactive astrocytes contributing to astrocyte scar formation [38]. In human, adult neurogenesis dropped sharply postnatally to undetectable levels in adults. Thus NSC transplantation is an ideal option to complement the necessary NPCs of required cell-fate. We found that the human NSCs could migrate extensively and tangentially along the white matter tracts, and enter the brain parenchyma radially, in concert with 2 previous studies by showing that the grafted human cells can migrate in the form of neural stem cell along the white matter tracts [1, 2]. But both studies only used Nestin as the NSC marker, leading to lots of debates about their conclusions.

Another important question is whether these migrating NSCs can differentiate into mature neurons and survive in the lesioned area. Although we failed to detect significant NeuN expression in human cells that reaching the distal areas, HuD/NeuN did express in the graft colonies, in cell cluster manner. One of the most important reasons is possibly that in the graft colony, the connections are readily formed in between the graft-derived neurons themselves, which are vital for the survival of neurons, especially in lack of the afferent inputs from the intact brain. As we known, neurons’ activity dictates their survival. The presynaptic neurons provide both depolarization inputs and anterograde neuronal trophic factors (e.g. BDNF) to the postsynaptic counterpart in an activity-dependent manner; meanwhile the postsynaptic cells might provide retrograde neurotrophic factors (e.g. NGF) to the presynaptic counterpart [39, 40]. In the migration pathway and the overlying cerebral cortex, where single neuroblasts or premature neurons might be eliminated gradually due to the low chance to form connections. This might represent a mechanism how the premature, dissociated or redundant neurons were removed during development and adult neurogenesis to maintain the homeostasis of the brain. Scenario might be different when the cells transplanted after brain injury, in which the denervated afferent and efferent host projections might provide an ideal environment for the graft survival and maturation. A recent study from Götz’ group gave an important support for this by showing that brain injury promoted initial integration of neuronal transplants and the excessive neurons at the lesion were pruned afterwards [9]. We have also shown that the human NPCs deposited in the infarction area provided potential innervating targets for host projection neurons and benefited to each other, and the physical exercises could further enhance their connections in an activity-dependent way [10]. However, before NPC therapy can enter the clinical trial, future work should be carried out to fully address two important questions, 1) how and to what extent the human long-term NPC grafts can be recruited by the brain damage, and if yes, 2) whether and how the brain damage can help the migratory NPCs survive and mature.

Last but not least, in case that the migratory NPCs gradually lost the neuronal differentiation potency, we yet have two choices to use them to achieve the cell therapy: 1) to convert them into neurons by inducible forced expression of ASCl1, alone or in combination with other pro-neuronal factors or small compounds [41-43]; 2) to use them as a potential source of cells for ***ex vivo*** gene therapy [44].

### The combination of *in vitro* neural derivatives and an *in vivo* physiological environment provides insights into the human neural development or adult neurogenesis

A big challenge to understand the human neural development and adult neurogenesis or neuro-regeneration is lack of access to brain tissue. Obviously, the ***in vivo*** ontogeny of human NPC grafts in adult rats we described here is quite different from both the *i**n vitro*** neural induction/differentiation and ***in vivo*** neural development and endogenous regeneration. Nevertheless, it could to some extent provide some insights into the human neural development or regeneration.

During the brain development in human and rodents, the GFAP up-regulation marks not only the transition of RGC differentiation competency from neurogenesis to gliogenesis (GFAP+/Nestin+: early RGC in human and late RGC in mice), but also the initial astrogliogenesis or the earliest fetal astrocytes (GFAP+/S100β+: at the beginning of mid-gestation in human and perinatally in mice) (for review, see [19, 20]. Meanwhile, ***in vitro*** neural induction and differentiation system has used GFAP as a reliable astrocyte marker, which was confirmed by co-expression of S100β, and validated via both ***in vitro*** and transplantation functional assays [15-17]. In our system, however, the majority of GFAP+ cells co-expressed a broad of NSC markers, like Nestin/Pax6/Sox2/HES5/ Vimentin/BLBP, but not the astrocyte maker S100β and GS, within a protracted period, showing great similarity to the human brain development, especially in terms of heterogeneity of NSC/NPCs. So the next important question is what the mechanism is that underlying the maintenance of NSC pool after transplantation in adult rats without a specialized NSC niche, like SVZ and SGZ.

Nowadays, brain organoids have been used as a powerful tool for studying of neural development and neural evolution of brain, as well some neuropsychiatric conditions [45], which stand at the interface of ***in vitro*** and ***in vivo***. However, brain organoids without a physiology microenvironment might have limitations to mature and form functional connections. A recent study from Sergiu Pașca’ group found that neurons in the human brain organoid grafts have a larger soma and more complicated dendrites, showing influence on rat behavior [46]. Mansour and colleagues also compared the ***in vitro*** organoid with the transplanted counterpart, and found that the organoid grafts have a much lower apoptotic level [47]. Both studies concluded that the combination of neural organoids and an ***in vivo*** physiological brain environment may benefit neural development as well as disease modeling.

Last but not least, the technical limitation of this study should be mentioned. Here we only used immunohistology to define the identity of human NSCs. Although we have tested as many markers and antibodies as we could, it was still hard to get a thorough view of the expression profile of the featured proteins in human cells, which however depends on the knowledge we have and good antibodies in hand. Recently, single cell sequencing has been widely used in this respect, due to its powerfulness in discovering new cell markers with high throughput and no need for antibodies. However, single cell sequencing also has its shortage, in that it cannot reflect the protein level, a fortiori neither the distribution of the markers to subcellular compartments. Thus in future studies, the combination of the single cell sequencing and histology immunostaining validation should be adopted to understand the underlying mechanisms, e.g. how the migratory human NSCs maintained, and how they respond to brain injury.

**In summary**, here we continuously trace the long-term cell fates and migration behaviors of the intra-striatal human NPC grafts. Together with accumulating evidences from other laboratories and ourselves, we propose that the grafted human NPCs might fulfill their cell therapy function via local and remote modes individually or in combination. Locally, they could differentiate into neurons with a relatively high efficiency, which then further mature and integrate into the host brain. Alternatively, the grafted human NPCs extensively migrate along the white matter tracts into the damaged brain parenchyma, where they might differentiate properly to replace the lost cells, or deliver neuroprotective and regenerative protein(s) to lesioned areas for ***ex vivo*** gene therapy. The local acting mode would be efficient for repairing local damages, while the remote mode should be the first choice for diffuse brain injury, e.g. Alzheimer’s disease. Further studies are needed to achieve these perspectives.

## Methods

All experiments followed the Guide for the Care and Use of Laboratory Animals of the National Institute of Health (8^th^ edition, 2011) and were approved by the Experimental Animal Ethics Committee at Guangzhou Institutes of Biomedicine and Health (GIBH), Chinese Academy of Sciences (IACUC NO. 2012008 and 2017056). All efforts were made to minimize animals used and any discomfort to the animals.

### Animals

Male Wistar or Sprague-Dawley (SD) rats aged 6–8 weeks were commercially provided by Charles River (Beijing, China) or the Animal Center of Southern Medical University (Guangzhou, China) and normally fed until an age of 8–10 weeks ready for surgery. Rats were maintained under SPF environment with a 12h/12h dark/light cycle. Upon transplantation, rats were anesthetized with 1.0–1.5% pentobarbital sodium (40–60 mg/kg body weight, i.p.). Atropine sulfate (Sigma, 0.05 mg/kg, s.c.) was used to reduce the sputum secretion. Cell transplantation was carried out on the stereotaxic device. Briefly, a minimal surgery above the injection site was performed and then after a thorough hemostasis, the NPCs were transplanted. After suture, rats were monitored to fully recover on a temperature-controlled blanket and returned to the cages with clean beddings. Antibiotics were applied daily in the first week after surgery.

### Neural induction and NPC transplantation

Human NPCs were induced from human embryonic stem cells (hESC Line H1 (passage 60–70), Wicell, Madison, WI, USA) as previously described by dual inhibition of SMAD signaling with minor modifications to get highly homogenous forebrain NPCs [4, 10, 21]. Briefly, 100% confluent hESCs were first induced to neuroepithelia monolayer within 8 days in N2B27 medium (DMEM/F12: neurobasal 1:1, 0.5% N2, 1% B27, 2mM Glutamax, 1 × NEAA, 5 μg/mL insulin, 2 μg/mL heparin) containing 5 μM SB431542 and 5 μM dorsomorphin. After passaging via Dispase digestion, cell patches were cultured in N2B27 medium for another 8 days. Upon the formation of typical rosette structures, NPCs were harvested by carefully picking of rosette structures and propagated on Matrigel-coated plate in N2B27 medium without bFGF and EGF. NPCs were routinely passaged via Accutase digestion about every 10 days until transplantation.

Upon transplantation, the NPCs were digested into single cells with Accutase and suspended in DMEM/F12 without any growth factors (7.5 × 10^4^ cells/ μL). 10× 10^4^ cells in total were microinjected in a single site (AP +1.0mm, LM 3.2mm, DV 4.0mm to Bregma). The needle was remained there for another 5 minutes before slowly withdrew.

### Immunohistology

At examining time points, subjects were euthanized with lethal pentobarbital sodium (200 mg/kg body weight, i.p.) and perfused transcardially with 0.9% NaCl followed by cold and fresh 4% paraformaldehyde in 0.1M phosphate buffer (pH 7.4). Brains were postfixed in 4% PFA overnight, and cryoprotected for 72 h in 30% sucrose at 4 °C. Brains were sliced coronally or sagittally to 40μm and processed for immunostaining. Free-floating immunohistochemistry was exploited to trace the cell fate of the human grafts. Sections were permeabilized with 1% Triton X-100 in PBS for 1 h, followed by incubating in blocking buffer (PBS containing 10% goat or donkey serum and 0.3% Triton X-100) for 1–2 h. Sections were then incubated in the primary antibodies diluted in blocking buffer at 4°C overnight and in Fluorescent secondary antibodies and DAPI for 1 h at room temperature. Sections were washed three times with PBS between steps, 15 min for each time on a shaker. Stained sections were mounted on a glass slide with an anti-fluorescence quenching mounting medium for further imaging and analysis.

Primary antibodies used in this study include: anti-human specific Nestin (**hNestin**, Abcam, ab6320, 1:500), anti-human specific Nestin (**hNestin**, Sigma, Abd69, 1:1000), anti-Sox2 (Abcam, ab97959, 1:500), anti-Pax6 (Covance Research, PRB-278P-100, 1:500), anti-Ki67 (Abcam, ab15580, 1:1000), anti-human specific nucleus(**hNA**, Millipore, MAB1281, 1:500), anti-BLBP (Cell Signaling,13347s, 1:2000), anti-Vimentin (Cell Signaling,5741,1:500), anti-Hes5 (Millipore, AB5708, 1:200), anti-human specific GFAP (**hGFAP**, TAKARA, **Stem123**, 1:1000), anti-AQP4 (Cell Signaling, 596785, 1:1000), anti-GS (Oasis biofarm,OB-PRB072-02, 1:100), anti-S100β (Abcam, ab52642, 1:200), anti-DCX (Cell Signaling, 4604, 1:500), anti-human specific cytoplasm (TAKARA, **Stem121**, 1:500), anti-TBR2 (Abcam, Ab23345, 1:200), anti-ASCL1 (Abcam, ab211327, 1:250), anti-NeuN (Millipore, ABN78,1:500), anti-HuD (Affinity, DF3251, 1:100), anti-NG2 (Cell Signaling, 43916, 1:250), anti-MBP (Millipore, AB980,1:500). Secondary antibodies include, Goat anti-Rabbit IgG (H+L), Alexa Fluor 647 (Invitrogen,A32733, 1:500), Goat anti-Rabbit IgG (H+L), Alexa Fluor 568 (Invitrogen, A11011,1:500), Goat anti-Rabbit IgG (H+L), Alexa Fluor 488 (Invitrogen,A11008, 1:500), Goat anti-Mouse IgG (H+L), Alexa Fluor 568 (Invitrogen, A11004, 1:500), Goat anti-Mouse IgG (H+L), Alexa Fluor 488 (Invitrogen, A11001, 1:500).

### Image analyzing and stereological techniques

For convenience, we grouped data from two near sampling time points into single assessment time point, e.g. 12 and 13 wpt, 16 and 17 wpt, 24 and 26 wpt, 38 and 41 wpt, respectively. For each marker at each assessment time point, 3 rats were included. 2-3 representative sections from each animal were analyzed, and 3 representative field centered on the target areas for each section were captured at 10X and 20X magnifications for quantification analysis. All images were captured on a Zeiss LSM800 confocal microscope and processed with Zen software and Adobe Illustrator (Adobe Systems, San Jose, CA).

For quantification of nucleus-located or cell-body stained markers, cell percentage analysis was used, e.g. Sox2/Pax6/ BLBP/Ki67 positive cells among human specific nucleus positive cells. The number of labeled cells in each representative field was stereologically estimated using the cell counter function of ImageJ software (NIH). Fiji automatic cell counting was used for field of low cell density, while cells were manually counted for images of high cell density using a Fiji plugin manual counter.

For markers that difficult to locate the individual cell body, percentages of overlapping fluorescence area were estimated, e.g. hGFAP among hNestin, hNestin among stem121 (human specific cytoplasm). The immunoreactivity areas for each marker were identified automatically by setting intensity threshold and quantified using ImageJ software.

All graphical data are presented as Means ± SEMs and were analyzed by using GraphPad Prism 6.0.

## Acknowledgements

This work was partially supported by the National Natural Science Foundation of China (81771356); the Science and Technology Programs of Guangzhou (201804020052, 202201020379, 201803040016); the Frontier and Key Technology Innovation Special Grant from the Department of Science and Technology of Guangdong Province (2020B1212060052).

## Author information

### Author Contributions

C.L. contributed to the experiments of neural induction, cell transplantation, immunostaining, data analysis and figure construction. A.L. did the immunostaining and image capture. Q.X. D.Z. W.M. and R.W. contributed to the animal surgery, the postoperative handling and animal perfusion. W.H. did the neural induction. H.S. did the postoperative handling and animal perfusion, and participated in double-blind experiments. X.H. contributed to technical support and double-blind experiments. D.Q. contributed to conception and design. X.Z. and X.W. contributed to data analysis and interpretation, and manuscript writing. Y.G. contributed to conception and design, data analysis and interpretation, and manuscript writing. All authors approved the submitted version.

## References

1. Englund, U., A. Bjorklund, and K. Wictorin, Migration patterns and phenotypic differentiation of long-term expanded human neural progenitor cells after transplantation into the adult rat brain. Brain Res Dev Brain Res, 2002. 134(1-2): p. 123–41.

2. Tabar, V., et al., Migration and differentiation of neural precursors derived from human embryonic stem cells in the rat brain. Nat Biotechnol, 2005. 23(5): p. 601–6.

3. Real, R., et al., In vivo modeling of human neuron dynamics and Down syndrome. Science, 2018. 362(6416).

4. Liu, C., et al., Hypoproliferative human neural progenitor cell xenografts survived extendedly in the brain of immunocompetent rats. Stem Cell Res Ther, 2021. 12(1): p. 376.

5. Liu, Y., et al., Medial ganglionic eminence-like cells derived from human embryonic stem cells correct learning and memory deficits. Nat Biotechnol, 2013. 31(5): p. 440–7.

6. Zhang, T., et al., Human Neural Stem Cells Reinforce Hippocampal Synaptic Network and Rescue Cognitive Deficits in a Mouse Model of Alzheimer’s Disease. Stem Cell Reports, 2019. 13(6): p. 1022–1037.

7. Palma-Tortosa, S., et al., Activity in grafted human iPS cell-derived cortical neurons integrated in stroke-injured rat brain regulates motor behavior. Proc Natl Acad Sci U S A, 2020. 117(16): p. 9094–9100.

8. Xiong, M., et al., Human Stem Cell-Derived Neurons Repair Circuits and Restore Neural Function. Cell Stem Cell, 2021. 28(1): p. 112–126 e6.

9. Grade, S., et al., Brain injury environment critically influences the connectivity of transplanted neurons. Sci Adv, 2022. 8(23): p. eabg9445.

10. Wu, R., et al., Physical exercise promotes integration of grafted cells and functional recovery in an acute stroke rat model. Stem Cell Reports, 2022. 17(2): p. 276–288.

11. Espuny-Camacho, I., et al., Pyramidal neurons derived from human pluripotent stem cells integrate efficiently into mouse brain circuits in vivo. Neuron, 2013. 77(3): p. 440–56.

12. Nagashima, F., et al., Novel and robust transplantation reveals the acquisition of polarized processes by cortical cells derived from mouse and human pluripotent stem cells. Stem Cells Dev, 2014. 23(18): p. 2129–42.

13. Kikuchi, T., et al., Human iPS cell-derived dopaminergic neurons function in a primate Parkinson’s disease model. Nature, 2017. 548(7669): p. 592-596.

14. Wang, Y.K., et al., Human Clinical-Grade Parthenogenetic ESC-Derived Dopaminergic Neurons Recover Locomotive Defects of Nonhuman Primate Models of Parkinson’s Disease. Stem Cell Reports, 2018. 11(1): p. 171–182.

15. Krencik, R., et al., Specification of transplantable astroglial subtypes from human pluripotent stem cells. Nat Biotechnol, 2011. 29(6): p. 528–34.

16. Windrem, M.S., et al., Human iPSC Glial Mouse Chimeras Reveal Glial Contributions to Schizophrenia. Cell Stem Cell, 2017. 21(2): p. 195–208 e6.

17. Canals, I., et al., Rapid and efficient induction of functional astrocytes from human pluripotent stem cells. Nat Methods, 2018. 15(9): p. 693–696.

18. Merkle, F.T., et al., Radial glia give rise to adult neural stem cells in the subventricular zone. Proc Natl Acad Sci U S A, 2004. 101(50): p. 17528–32.

19. Gotz, M. and Y.A. Barde, Radial glial cells defined and major intermediates between embryonic stem cells and CNS neurons. Neuron, 2005. 46(3): p. 369–72.

20. Sardar, D., et al., Mechanisms of astrocyte development., in Patterning and cell type specification in the developing CNS and PNS, J. Rubernstein and P. Rakic, Editors. 2020, Elsevier Inc. p. 807-827.

21. Xing, Q., et al., Retrograde monosynaptic tracing through an engineered human embryonic stem cell line reveals synaptic inputs from host neurons to grafted cells. Cell Regen, 2019. 8(1): p. 1–8.

22. Gotz, M., M. Nakafuku, and D. Petrik, Neurogenesis in the Developing and Adult Brain-Similarities and Key Differences. Cold Spring Harb Perspect Biol, 2016. 8(7).

23. Fu, Y., et al., Heterogeneity of glial progenitor cells during the neurogenesis-to-gliogenesis switch in the developing human cerebral cortex. Cell Rep, 2021. 34(9): p. 108788.

24. Ziffra, R.S., et al., Single-cell epigenomics reveals mechanisms of human cortical development. Nature, 2021. 598(7879): p. 205-213.

25. Feng, L., M.E. Hatten, and N. Heintz, Brain lipid-binding protein (BLBP): a novel signaling system in the developing mammalian CNS. Neuron, 1994. 12(4): p. 895–908.

26. Basak, O. and V. Taylor, Identification of self-replicating multipotent progenitors in the embryonic nervous system by high Notch activity and Hes5 expression. Eur J Neurosci, 2007. 25(4): p. 1006–22.

27. Edri, R., et al., Analysing human neural stem cell ontogeny by consecutive isolation of Notch active neural progenitors. Nat Commun, 2015. 6: p. 6500.

28. Holst, C.B., et al., Astrogliogenesis in human fetal brain: complex spatiotemporal immunoreactivity patterns of GFAP, S100, AQP4 and YKL-40. J Anat, 2019. 235(3): p. 590-615.

29. Habbas, S., et al., Neuroinflammatory TNFalpha Impairs Memory via Astrocyte Signaling. Cell, 2015. 163(7): p. 1730–41.

30. Eelen, G., et al., Role of glutamine synthetase in angiogenesis beyond glutamine synthesis. Nature, 2018. 561(7721): p. 63-69.

31. Donato, R., et al., Functions of S100 proteins. Curr Mol Med, 2013. 13(1): p. 24–57.

32. Fallier-Becker, P., et al., Onset of aquaporin-4 expression in the developing mouse brain. Int J Dev Neurosci, 2014. 36: p. 81–9.

33. Castaneyra-Ruiz, L., et al., AQP4 labels a subpopulation of white matter-dependent glial radial cells affected by pediatric hydrocephalus, and its expression increased in glial microvesicles released to the cerebrospinal fluid in obstructive hydrocephalus. Acta Neuropathol Commun, 2022. 10(1): p. 41.

34. Herzog, C., et al., Rapid clearance of cellular debris by microglia limits secondary neuronal cell death after brain injury in vivo. Development, 2019. 146(9).

35. Kriks, S., et al., Dopamine neurons derived from human ES cells efficiently engraft in animal models of Parkinson’s disease. Nature, 2011. 480(7378): p. 547-51.

36. Song, B., et al., Human autologous iPSC-derived dopaminergic progenitors restore motor function in Parkinson’s disease models. J Clin Invest, 2020. 130(2): p. 904–920.

37. Gritti, A., et al., Multipotent neural stem cells reside into the rostral extension and olfactory bulb of adult rodents. J Neurosci, 2002. 22(2): p. 437–45.

38. Faiz, M., et al., Adult Neural Stem Cells from the Subventricular Zone Give Rise to Reactive Astrocytes in the Cortex after Stroke. Cell Stem Cell, 2015. 17(5): p. 624–34.

39. Hefti, F., Nerve growth factor promotes survival of septal cholinergic neurons after fimbrial transections. J Neurosci, 1986. 6(8): p. 2155–62.

40. Ghosh, A., J. Carnahan, and M.E. Greenberg, Requirement for BDNF in activity-dependent survival of cortical neurons. Science, 1994. 263(5153): p. 1618-23.

41. Pang, Z.P., et al., Induction of human neuronal cells by defined transcription factors. Nature, 2011. 476(7359): p. 220-3.

42. Guo, Z., et al., In vivo direct reprogramming of reactive glial cells into functional neurons after brain injury and in an Alzheimer’s disease model. Cell Stem Cell, 2014. 14(2): p. 188–202.

43. He, S., et al., Reprogramming somatic cells to cells with neuronal characteristics by defined medium both in vitro and in vivo. Cell Regen, 2015. 4: p. 12.

44. Emborg, M.E., et al., GDNF-secreting human neural progenitor cells increase tyrosine hydroxylase and VMAT2 expression in MPTP-treated cynomolgus monkeys. Cell Transplant, 2008. 17(4): p. 383–95.

45. Lu, K., et al., Depressive patient-derived GABA interneurons reveal abnormal neural activity associated with HTR2C. EMBO Mol Med, 2023. 15(1): p. e16364.

46. Revah, O., et al., Maturation and circuit integration of transplanted human cortical organoids. Nature, 2022. 610(7931): p. 319-326.

47. Mansour, A.A., et al., An in vivo model of functional and vascularized human brain organoids. Nat Biotechnol, 2018. 36(5): p. 432–441.

